# PHF1 compartmentalizes PRC2 at target loci via phase separation

**DOI:** 10.1101/2022.10.20.513034

**Authors:** Genzhe Lu, Pilong Li

## Abstract

Polycomb repressive complex 2 (PRC2) is central to polycomb repression as it trimethylates lysine 27 on histone H3 (H3K27me3). How PRC2 is recruited to its targets to deposit H3K27me3 remains an open question. Polycomb-like (PCL) proteins, a group of conserved PRC2 accessory proteins, direct PRC2 to its targets. In this report, we demonstrate that a PCL protein named PHF1 forms phase-separated condensates at H3K27me3 loci that recruit PRC2. Combining cellular observation and biochemical reconstitution, we show that the N-terminal domains of PHF1 cooperatively mediate target recognition, the chromo-like domain recruits PRC2, and the intrinsically disordered region (IDR) drives phase separation. Moreover, we reveal that the condensates compartmentalize PRC2, DNA, and nucleosome arrays by phase separation. Luciferase reporter assays confirm that PHF1 phase separation promotes transcription repression, further supporting a role of the condensates in polycomb repression. Based on our findings, we propose that these condensates create favorable microenvironments at the target loci for PRC2 to function.

## Introduction

As the sole methyltransferase in mammalian cells responsible for trimethylation of lysine 27 on histone H3 (H3K27me3), polycomb repressive complex 2 (PRC2) is essential for gene repression, and its malfunction results in developmental disorders^1–3^. The PRC2 core complex is composed of Suz12, EZH1/2, EED, and Rbbp4/7, among which Suz12 interacts with all subunits to organize the complex and EZH1/2 possesses the methyltransferase activity^4,5^. Based on the accessory proteins associated with the core complex, PRC2 is divided into two biochemically distinct subtypes – PRC2.1 and PRC2.2. PRC2.1 is characterized by the association of a polycomb-like (PCL) protein (PHF1, MTF2, or PHF19) and EPOP or PALI1/2, whereas PRC2.2 contains AEBP2 and JARID2^1,6^.

Despite intensive investigations, it remains unclear how PRC2 is recruited to its targets to deposit H3K27me3^7^. PCL proteins, which are components of PRC2.1, direct PRC2 to its targets for H3K27me3 deposition^8–12^. A recent study showed that H3K27me3 was first deposited at “nucleation sites”, and then spread to distal regions^13^. The depletion of MTF2, one of the PCL proteins, causes a severe delay in this process in mouse embryonic cells^13^. Upon the depletion of all PCL proteins, the H3K27me3 level decreases, whereas the H3K27me1 and me2 levels increase^12,14^. This is in line with the fact that the PRC2 core complex efficiently catalyzes H3K27me1/2, but not H3K27me3, whereas PCL proteins enhance the H3K27me3 activity, partly by prolonging the residence time of PRC2 on chromatin^15–18^. Of note, only H3K27me3, but not H3K27me1/2, underlies cell type-specific gene repression^19^. Functionally, PCL proteins are involved in hematopoiesis, spermatogenesis, DNA repair, prostate cancer, and embryonic stem cell differentiation^10,20–24^.

PCL proteins share a similar domain topology (Figure 1A). PCL proteins possess a Tudor domain, two PHD finger domains (PHD1 and 2), and an extended homologous (EH) domain in the N-terminal moiety. Notably, the EH domain specifically recognizes unmethylated CpG islands, which coincide with many polycomb targets^16,25–27^. The chromo-like domain at the C-terminus of PCL proteins binds Suz12, thereby connecting PCL proteins to the PRC2 core complex^28^. Interestingly, the association of the chromo-like domain to the PRC2 core complex elicits the dimerization of PRC2, and consequently promotes CpG DNA binding *in vitro*^28^. Additionally, there is a positively charged intrinsically disordered region (IDR) between the EH domain and the chromo-like domain with unknown functions (Figures 1A and S1).

**Figure 1.**
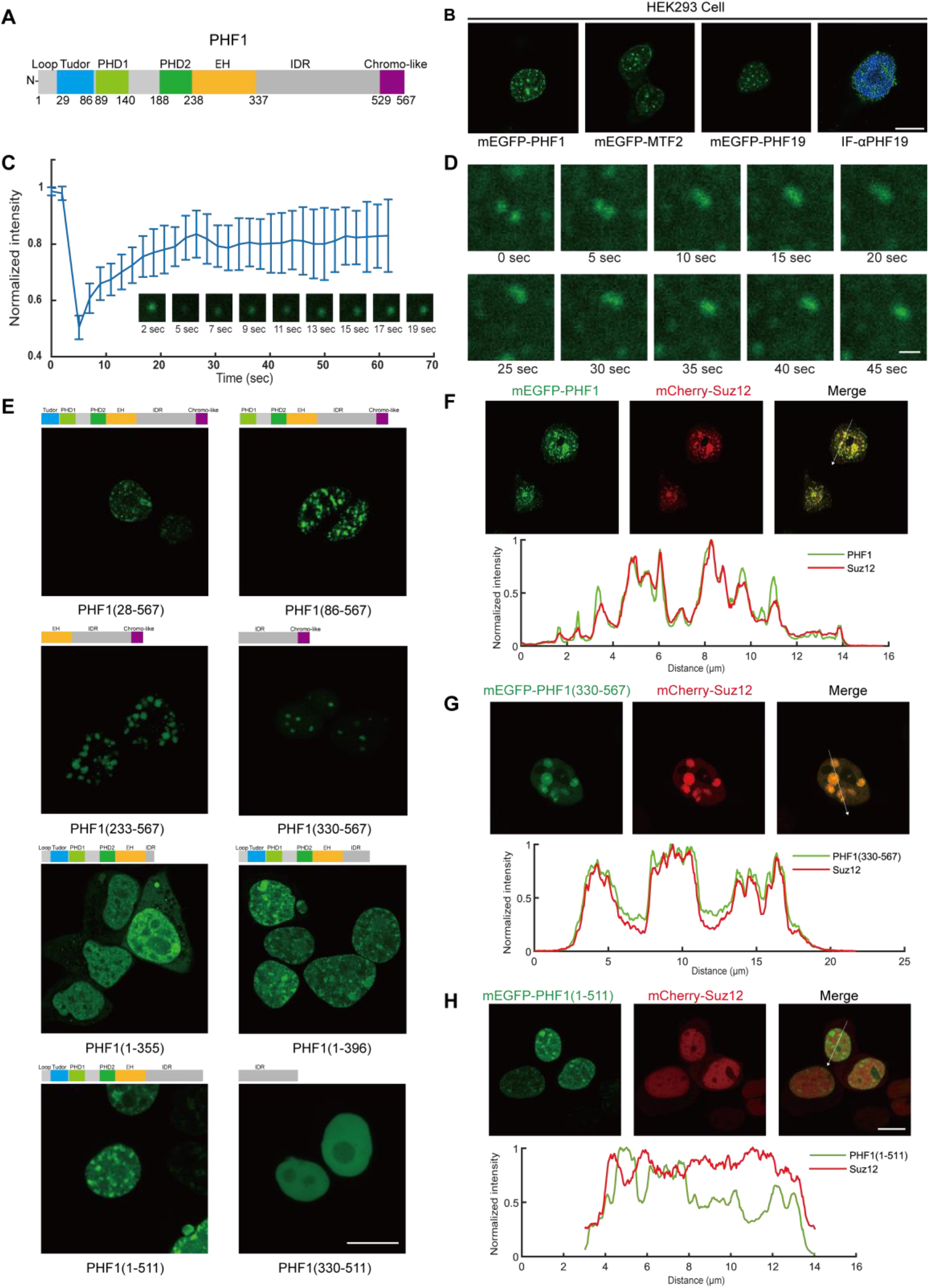
PHF1 forms biomolecular condensates that recruit Suz12 *in vivo*. **A** Domain topology of PHF1. **B** Confocal images of mEGFP-PHF1, mEGFP-MTF2, and mEGFP-PHF19 in living cells, and immunofluorescence staining of PHF19 (IF-αPHF19) in HEK293 cells. Nuclei were stained by DAPI. Scale bar, 10 μm. **C** Fluorescence recovery after photobleaching (FRAP) analysis of PHF1 puncta in HEK293 cells (n=7). **D** Fusion of PHF1 puncta in HEK293 cells. Scale bar, 0.5 μm. **E** Confocal images of the indicated mEGFP-tagged PHF1 truncations after transfection into HEK293 cells. Scale bar, 10 μm. **F to H** Co-transfection of PHF1 variants and Suz12. In each subfigure, the upper panel shows confocal images of HEK293 cells cotransfected with mCherry-Suz12 and mEGFP-PHF1 (**F**), -PHF1(330-567) (**G**), or -PHF1(1-511) (**H**); and the lower panel shows line-scan analyses of fluorescence signals along the arrow line in the Merge image. Scale bar, 10 μm.

In this report, we show that IDR of PHF1 phase-separates to form condensates that recruit PRC2 to its target sites. Liquid-liquid phase separation (LLPS) has emerged as a novel mechanism whereby large biomolecular assemblies form for precise spatiotemporal regulation of multiple biological processes^29,30^. We propose that PCL proteins recruit PRC2 to execute gene repression by compartmentalization at the target loci rather than via simple positioning.

## Results

### PCL proteins form biomolecular condensates in vivo

To examine the intracellular distribution of PCL proteins, we performed *in vivo* imaging of PCL proteins. We transiently transfected mEGFP-tagged PHF1, MTF2, and PHF19 into HEK293 cells or HeLa cells, and performed confocal live-cell imaging. Upon overexpression, PCL proteins formed punctum-like condensates in living cells (Figures 1B and S2). To corroborate the condensate formation in the native state, we performed immunofluorescence staining of endogenous PHF19 in HEK293 cells and HeLa cells. We chose PHF19 because of its relatively high expression levels in these cell lines. Endogenous PHF19 was present in small but distinct nuclear puncta (Figures 1B and S2). The fast fluorescence recovery after photobleaching (FRAP) and the rapid fusion of PHF1 puncta indicated the dynamic nature of the condensates (Figures 1C and 1D). These observations suggest that PCL proteins have the potential to form biomolecular condensates *in vivo*, which is suggestive of a liquid-liquid phase separation mechanism.

### The IDR of PHF1 is required for condensate formation

We chose PHF1 as a representative PCL protein to further investigate the mechanism underlying PCL condensate formation. In addition to the structured domains (Figure 1A), there is a positively charged IDR between the EH domain and the chromo-like domain of PCL proteins (Figures 1A and S1). We constructed a series of mEGFP-tagged PHF1 truncations to map the essential regions responsible for condensate formation. We found that the deletions of the N-terminal loop (PHF1(28-567)), the Tudor domain (PHF1 (86-567)), the N-terminal moiety except for the EH domain (PHF1(233-567)), and the entire N-terminal moiety (PHF1(330-567)) of PHF1 did not affect the punctum-like morphology in living cells (Figure 1E). In contrast, deleting the IDR of PHF1 abolished the formation of puncta (Figure 1E; PHF1(1-355)). Interestingly, the puncta seemed to reappear as longer IDR segments were preserved in the truncations (Figure 1E; compare PHF1(1-355), PHF1(1-396), and PHF1(1-511)). Unexpectedly, the IDR segment alone was not able to form puncta in living cells (Figure 1E, PHF1(330-511)), which implies that additional mechanisms aid the formation of condensates in cells. In summary, these results demonstrate that the IDR of PHF1 is required for the formation of PHF1 condensates *in vivo*.

### Suz12 is recruited into PHF1 condensates by the chromo-like domain

Among the multiple functions of PCL proteins, recruiting and localizing PRC2 is an important one^10,12,14,25^. Therefore, we next sought to examine the interplay between PHF1 and Suz12, which is the organizing center of PRC2. We co-transfected mCherry-tagged Suz12 and mEGFP-tagged PHF1 variants into HEK293 cells (Figures 1F–1H). In the cells transfected with full-length PHF1 or PHF1(330-567), Suz12 was recruited into the condensates (Figures 1F and 1G). In contrast, Suz12 was not enriched in condensates formed by PHF1(1-511), which lacked the chromo-like domain (Figure 1H). This is consistent with a previous report showing that the chromo-like domain binds the C2 domain of Suz12^28^. In conclusion, the chromo-like domain of PHF1 mediates the recruitment of PRC2 to the condensates by specifically binding Suz12.

### N-terminal domains of PHF1 cooperatively contribute to target recognition

PCL proteins colocalize with H3K27me3, as evidenced by ChIP-seq data and imaging data^12,14,21,31^. To corroborate this, we performed immunofluorescence staining of H3K27me3 in HEK293 cells transfected with PHF1. As expected, PHF1 condensates colocalized with H3K27me3 loci (Figure 2A and S3). Biochemical studies have shown that several domains in the N-terminal moiety of PCL proteins recognize certain markers on chromatin; specifically, the Tudor domain recognizes H3K36me3, the PHD1 domain recognizes H4R3 symmetric dimethylation, and the EH domain recognizes unmethylated CpG islands^8–11,25,26,32^. To dissect how these domains contribute to PHF1 target recognition, we introduced loss-of-function point mutations into the Tudor domain (Y47A), the PHD1 domain (H115A), and the EH domain (K323A) of PHF1^9,26,32^, and examined the colocalization of condensates with H3K27me3 loci. We found that the condensates failed to colocalize with H3K27me3 loci when any one of the aforementioned domains was mutated (Figures 2B–2D and S3). Additionally, a PHF1 truncation with an intact N-terminal moiety (PHF1(1-396)) was sufficient to mediate colocalization with H3K27me3, whereas the C-terminal moiety of PHF1 (PHF1(330-567)) was not (Figures 2E, 2F, and S3). In conclusion, the Tudor domain, the PHD1 domain, and the EH domain cooperatively contribute to PHF1 target recognition.

**Figure 2.**
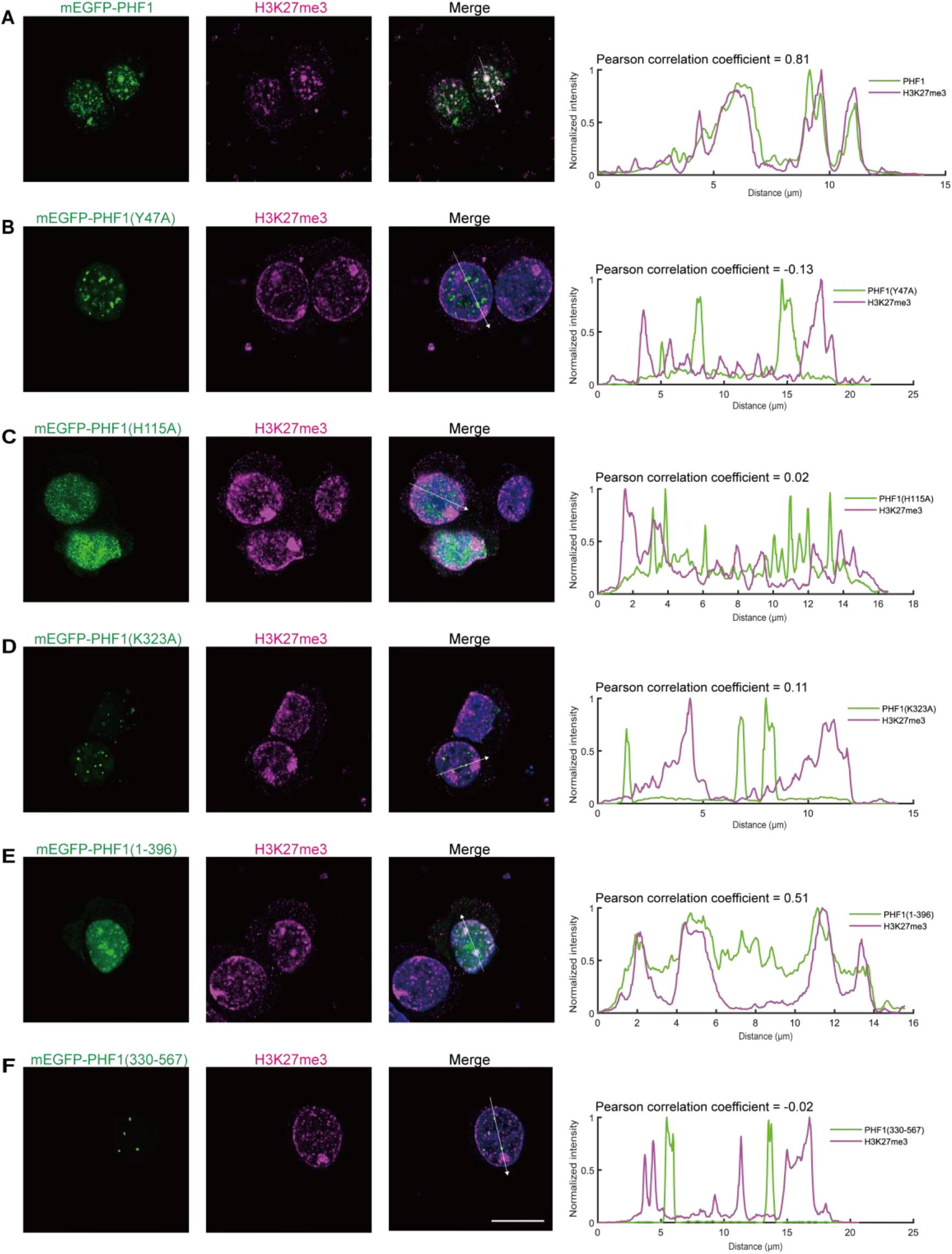
The Tudor domain, PHD1 domain, and EH domain cooperatively contribute to the localization of PHF1 to PRC2 targets. **A to F** Immunofluorescence staining of H3K27me3 in HEK 293 cells transfected with mEGFP-tagged PHF1 (**A**), PHF1(Y47A) (**B**), PHF1(H115A) (**C**), PHF1(K323A) (**D**), PHF1(1-396) (**E**), and PHF1(330-567) (**F**). The rightmost panel in each subfigure shows the line scan analysis of fluorescence signals along the arrow line in the Merge image. The Pearson correlation coefficient for colocalization of PHF1 variants and H3K27me3 is shown above each line scan trace. Scale bar, 10 μm. See also Figure S3 for Pearson correlation analyses of more cells.

Combining the data from the regional mapping and Suz12 recruitment experiments, we conclude here that the N-terminal moiety of PHF1 is responsible for target recognition, the IDR of PHF1 is required for condensate formation, and the chromo-like domain is responsible for PRC2 recruitment *in vivo*. By compartmentalizing PRC2 at the target loci, these condensates may promote polycomb repression.

### PHF1-IDR undergoes phase separation in vitro

To test the hypothesis that the IDR of PHF1 mediates the formation of condensates by phase separation, we purified mEGFP-tagged PHF1(330-511) (referred to as PHF1-IDR hereafter) and performed *in vitro* phase separation assays. We found that PHF1-IDR underwent phase separation under low-salt conditions without crowding agents (Figure 3A). FRAP analysis indicated that the condensates were dynamic (Figure 3B). Phase diagram analysis showed that condensate formation was dependent on protein concentrations and ionic strengths (Figure 3C). High ionic strengths disrupted the condensates, indicating that electrostatic interactions between PHF1-IDR molecules contribute to the formation of condensates (Figures 3C and S1). Although phase-separated condensates of PHF1-IDR were not observed in living cells, we have demonstrated above that the IDR was required for PHF1 condensate formation (Figure 1E). Therefore, we conclude that the formation of PHF1 condensates is largely driven by the phase separation capability of PHF1-IDR, which may be further potentiated by other factors such as chromatin.

**Figure 3.**
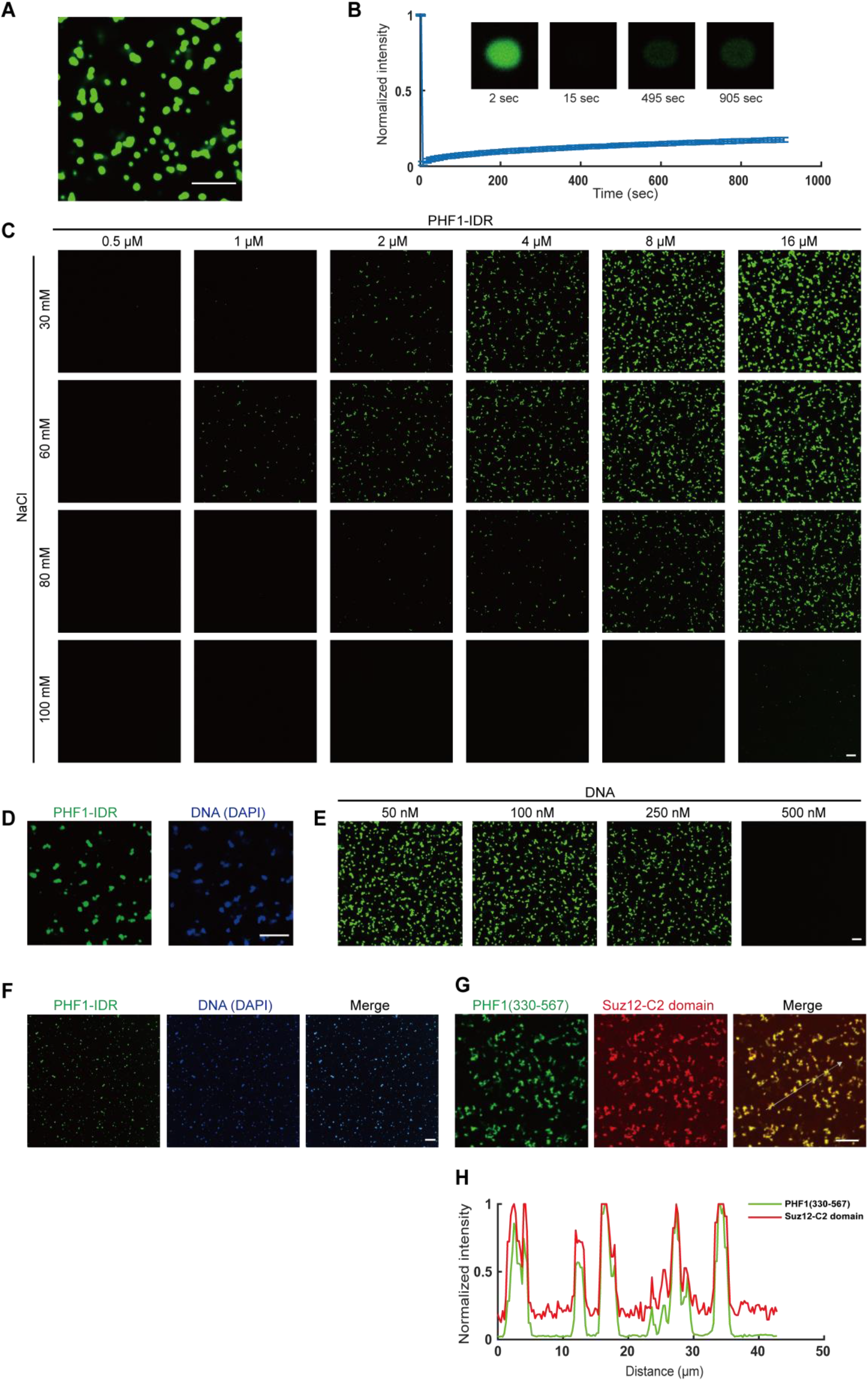
PHF1-IDR undergoes phase separation *in vitro*. **A** Representative image of PHF1-IDR condensates. **B** FRAP analysis of PHF1-IDR condensates (n=12). **C** Phase diagram of PHF1-IDR concentrations with respect to salt concentrations. **D** Representative image of cocondensates of 16 μM PHF1-IDR with 100 nM 72 bp DNA. **E** Impact of DNA on formation of PHF1-IDR condensates. See also figure S5. **F** Compaction of 185 nM H3K27me3 nucleosome arrays by 2 μM PHF1-IDR at 100 mM NaCl. **G** Recruitment of the Suz12-C2 domain into PHF1(330-567) condensates. **H** Line scan analysis of fluorescence signals along the arrow line in the Merge image of G. All scale bars, 10 μm.

### PHF1 compartmentalizes its nuclear partners by phase separation

PHF1 is surrounded by DNA and nucleosomes at its target loci. To investigate the interplay between PHF1 condensates and DNA or nucleosome arrays, we performed a series of co-condensation assays. We found that DNA was able to partition into the PHF1-IDR condensates (Figure 3D). We reasoned that the partitioning of DNA into the condensates was driven by sequence-nonspecific electrostatic interactions between the positively charged PHF1-IDR and the negatively charged DNA backbone. Consistent with this idea, increasing DNA concentrations disrupted the condensates, presumably resulting from disruption of the charge balance^33^ (Figures 3E and S5). Recent studies have shown that nucleosome arrays (NA) can act as a scaffold for phase separation by providing multivalency^34–36^. To examine how PHF1-IDR affects chromatin compaction, we co-incubated PHF1-IDR and 12X H3K27me3-modified nucleosome arrays (12X H3K27me3 NA) under conditions where PHF1-IDR and nucleosome arrays *per se* did not phase separate (100 mM NaCl, Figures 3C and S6). We found that the 12X H3K27me3 NA co-phase separated with PHF1-IDR in this system (Figure 3F), which indicates that NA indeed enhanced phase separation of PHF-IDR. Such a co-phase separation scenario might be more physiologically relevant owing to the participation of nucleosome arrays and lower protein concentrations. We reasoned that the binding of the PHF1 N-terminal moiety to target loci results in local nucleation of chromatin and PHF1-IDR for co-phase separation. This is consistent with the finding that PHF1(1-511) forms biomolecular condensates in cells whereas PHF1-IDR does not (Figures 1E).

To corroborate PRC2 recruitment by PHF1 condensates *in vitro*, we purified mEGFP-tagged PHF1(330-567) and mCherry-tagged Suz12-C2 domain and performed a recruitment assay. The result showed that the Suz12-C2 domain was recruited into PHF1(330-567) condensates (Figures 3G and 3H). In summary, these *in vitro* reconstitution data indicate that PHF1 is capable of enriching PRC2 and compacting chromatin by phase separation. Since nucleosomes are the substrate of PRC2, we speculate that such compartmentalization and compaction create favorable microenvironments, at least by substantially increasing the local concentrations of both, for PRC2 to deposit H3K27me3.

### Phase separation by PHF1-IDR promotes transcription repression in cells

We were curious about whether the compartmentalization of PRC2 at the target loci by phase-separated PHF1 condensates could promote transcription repression. To this end, we designed a luciferase reporter system in which 9 tandem GAL4-binding sites were cloned upstream of a herpes simplex virus thymidine kinase (HSV-TK) promoter followed by a firefly luciferase gene (Figure 4A). We expressed a series of GAL4-fused PHF1 variants to examine their effects on luciferase transcription (Figure S7). GAL4-PHF1-IDR did not cause transcription repression in comparison with GAL4 alone, which indicates that the recruitment of PRC2 to the target by the chromo-like domain is required for transcription repression (Figure S7). Transcription repression by full-length PHF1 (GAL4-PHF1FL) was not as strong as GAL4-PHF1-IDR-chromo-like and GAL4-PHF1-chromo-like. This result is presumably explained by the competition between the luciferase reporter plasmid and endogenous target loci for PHF1 binding (Figure S7).

**Figure 4.**
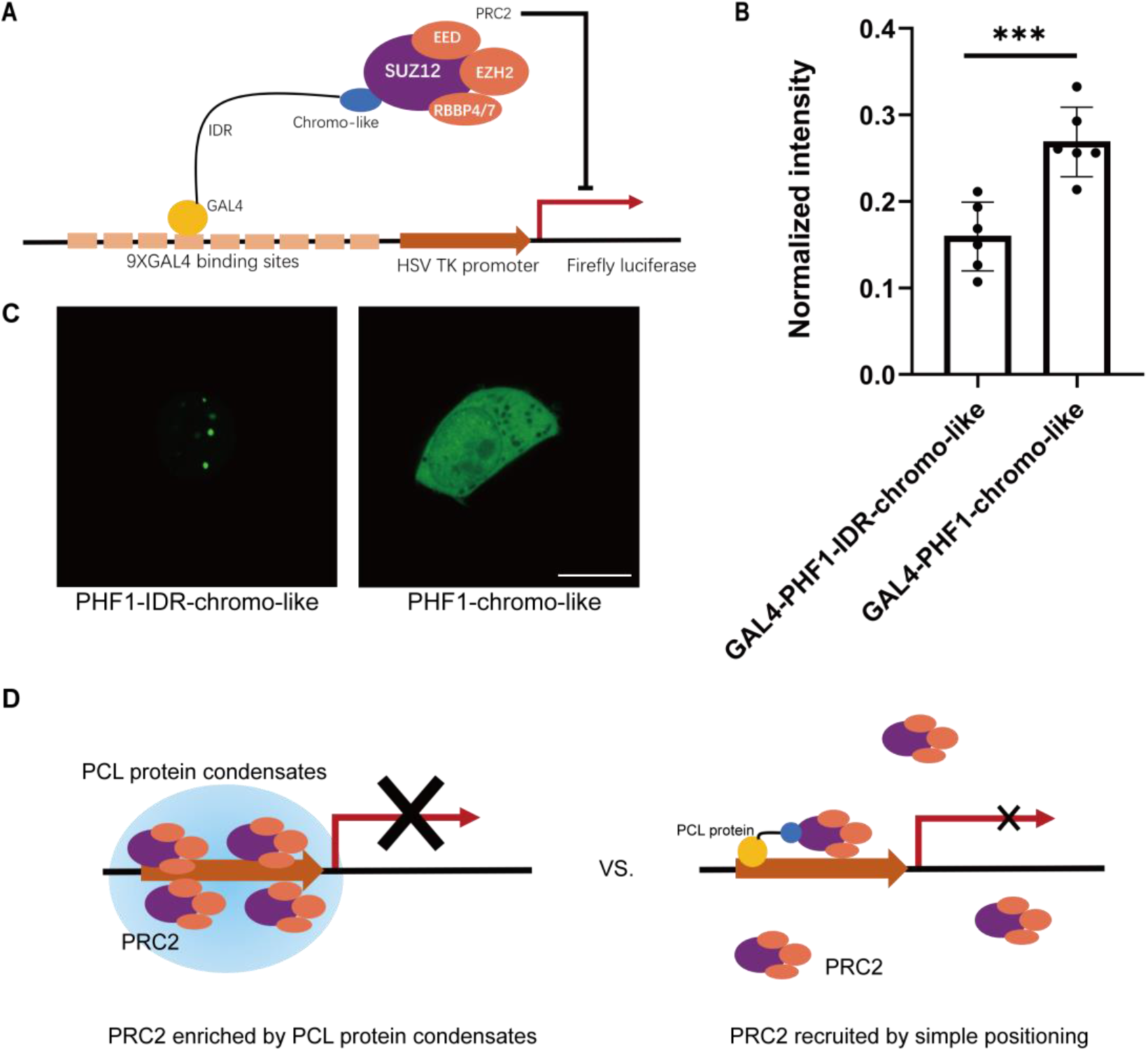
Phase separation by PHF1-IDR promotes transcription repression. **A** Schematic illustration of the luciferase reporter assay. **B** Luciferase reporter assay results for GAL4-PHF1-IDR-chromo-like and GAL4-PHF1-chromo-like (n=6). The luminescence intensities were normalized (see Methods). Unpaired Student’s t-test was used to compare the two groups (***, p<0.001). **C** Representative confocal images of mEGFP-tagged PHF1-IDR-chromo-like (residues 330-567) and PHF1-chromo-like (residues 512-567). Scale bar, 10 μm. **D** A model summarizing the results of this work. PCL proteins direct and enrich PRC2 at its targets by phase separation to promote gene repression.

Among these different PHF1 constructs, we noted that GAL4-PHF1-IDR-chromo-like caused significantly stronger transcription repression than GAL4-PHF1-chromo-like (Figure 4B). We found that mEGFP-tagged PHF1-IDR-chromo-like formed phase-separated condensates in living cells, while mEGFP-tagged PHF1-chromo-like did not (Figure 4C). Therefore, we reasoned that GAL4-PHF1-chromo-like recruits PRC2 to the GAL4-binding sites by simple positioning to repress transcription. In contrast, by forming phase-separated condensates, GAL4-PHF1-IDR-chromo-like provides an extra layer of enrichment of PRC2 and DNA, which leads to stronger transcription repression (Figure 4D). Based on these results, we conclude that phase separation by PHF1 promotes the transcription repression activity of PRC2.

## Discussion

LLPS is involved in numerous biological processes^29,30^. In this work, we demonstrated that PHF1 forms phase-separated condensates both *in vivo* and *in vitro*. We also found that PHF1 condensates colocalize with H3K27me3 loci, recruit and enrich Suz12, compact DNA and nucleosome arrays, and promote transcription repression. Mechanistically, the N-terminal moiety of PHF1 mediates target recognition, the IDR drives phase separation, and the chromo-like domain recruits PRC2. These lines of evidence indicate that PCL proteins compartmentalize PRC2 at target loci by phase separation.

Unlike AEBP2 and JARID2, which activate PRC2 allosterically, PCL proteins activate PRC2 by directing PRC2 to its targets and prolonging the residence time on chromatin^4,16,17,37^. Although PCL proteins can form stoichiometric complexes with the PRC2 core complex *in vitro*^16,38^, our results suggest that PCL proteins have a tendency to self-assemble into phase-separated condensates, which is distinct from the simple positioning mechanism of stoichiometric complexes (Figure 4D). We posit that there is an equilibrium between PCL proteins in the phase-separated condensates and in stoichiometric PRC2.1 complexes, through which the cell can regulate the dynamics of PRC2.1.

It is believed that PRC1 carries out compaction of facultative heterochromatin, likely by phase separation^1,39–45^. However, several lines of evidence suggest that PRC2 is likewise able to participate in chromatin compaction^46–49^. Our results suggest that a group of PRC2 accessory proteins is also capable of compacting chromatin by phase separation. By compacting chromatin and compartmentalizing PRC2 at the target loci, PCL condensates create favorable microenvironments for PRC2 to function. Since PCL proteins are important for *de novo* establishment of H3K27me3, it is plausible that PCL condensates act as hubs for H3K27me3 nucleation and spreading^13^. The polycomb-mediated chromatin compaction modes mentioned above are not mutually exclusive. On the contrary, they may contribute to polycomb repression concomitantly. Further investigations are required to determine whether, and if so how, these modes of compaction result in transcription repression.

## Methods

### Molecular cloning

Human PHF1, MTF2, PHF19, and Suz12 cDNAs were cloned into pCDNA3.1 eukaryotic expression vectors with mEGFP or mCherry at the N-terminus. PHF1 truncations were constructed by PCR. PHF1 point-mutated variants were generated using a point mutation kit (TransGen Biotech).

### Cell culture and transfection

HEK293 cells and HeLa cells were cultured in high glucose DMEM medium (Hyclone) supplemented with 10% fetal bovine serum (FBS), 100 μg/mL penicillin, 100 μg/mL streptomycin, and 2 mM L-glutamine at 37 °C, 5% CO2. For transfection, the cells were seeded in a 4-chamber glass-bottom dish (Invitrogen) and transfected with 1 μg plasmids using 2 μg PEI as the transfection reagent.

### Immunofluorescence staining

Cells were fixed for 15 minutes using 4% paraformaldehyde fix solution (Beyotime), washed with PBS, permeabilized with enhanced immunostaining permeabilization buffer (Beyotime), and blocked with immunostaining blocking buffer (Beyotime). Then the cells were incubated with primary antibodies at 4 °C overnight, washed with PBS, and incubated with secondary antibodies at room temperature for 2 hours followed by a PBS wash. The sample was subjected to imaging after adding antifade mounting medium with DAPI (Beyotime).

### Protein expression and purification

mEGFP-PHF1-IDR and mEGFP-PHF1(330-567) were cloned into pMAL (Synbio Technologies) bacterial expression vectors, with a his6 tag at the C-terminus. The N-terminal MBP tag was followed by a TEV protease cleavage site and then mEGFP. Rosetta(DE3) cells (KT Health) transformed with expression vectors were cultured in LB medium at 37 °C to O.D. = 1.0, and then induced with 0.8 mM IPTG at 16 °C overnight. The bacterial cells were harvested by centrifugation and stored at −80 °C. For purification, the cell pellets were thawed in lysis buffer (40 mM Tris, 500 mM NaCl, 5% glycerol, 30 mM imidazole, 2 mM BME, 1 mM PMSF, pH = 7.5), lysed by high-pressure cracking and sonication, and clarified by centrifugation at 18,000 rpm for 40 minutes. The supernatant was passed through a Ni-NTA column and washed extensively with wash buffer (40 mM Tris, 1 M NaCl, 5% glycerol, 40 mM imidazole, pH = 7.5). Then the protein was eluted by elution buffer (40 mM Tris, 100 mM NaCl, 5% glycerol, 500 mM imidazole, pH = 7.5). TEV protease was added to the eluate at 4 °C overnight to cleave off the MBP tag. The protein was loaded onto an SP column (GE Healthcare) and eluted with a linear NaCl gradient from 10 mM to 800 mM. The MBP tag had no affinity for the SP resin, and was therefore removed in this step. The peak fractions were pooled, concentrated, and passed through a Superdex 200 Increase 10/300 GL (GE Healthcare) gel filtration column using SEC running buffer (50 mM HEPES, 300 mM NaCl, 5% glycerol, pH = 7.5). The peak fractions were pooled, concentrated, and snap-frozen in liquid N2.

mCherry-Suz12-C2 domain (residues 146-363) was cloned into a PGEX-4T 1 vector and expressed, lysed, and clarified as described above. The lysate was passed through a GST column. The protein was eluted by on-column TEV protease cleavage.

### *In vitro* phase separation assay

The total volume of a phase separation system was 20 μL. Each phase separation system was prepared in a 200 μL PCR tube with NaCl-free dilution buffer (50 mM HEPES, 5% glycerol, pH = 7.5), and then transferred into a glass-bottom 384-well plate (Cellvis) for imaging. The 72 bp dsDNA (forward strand sequence: 5’-GCCACCGGTGGCTTCTTCTAGCCACCGGTGGCTTCTTCTAGCCACCGGTGGCTTCTTCTAGCCACCGGTGGC-3’) was synthesized and annealed in annealing buffer (40 mM Tris, 50 mM NaCl, 5% glycerol). Nucleosome arrays, which were prepared as described previously^35,50^, were a gift from Wenjing Sun.

### Luciferase reporter assay

The luciferase reporter assay was performed as previously described^51^. 9X GAL4-binding sites and an HSV TK promoter were cloned upstream of the firefly luciferase gene in the pGL4.27 vector (Promega, Cat. E8451), yielding the luciferase reporter vector. GAL4-PHF1 variants were cloned into pCDA 3.1 eukaryotic expression vectors. 300 ng of luciferase reporter vector, 300 ng of the protein expression vector, and 6 ng of *Renilla* luciferase vector (Promega, Cat. E6921) were co-transfected into HEK 293 cells. Luciferase luminescence intensities were measured 24 hours after transfection using the Dual-Luciferase^®^ Reporter Assay System (Promega). The luminescence intensities of firefly luciferase were normalized to that of *Renilla* luciferase. Subsequently, the luminescence intensities were normalized to the mock group. 6 biological replicates were performed.

### Data analysis

Images were processed in a Nikon-Elements image processing platform. All error bars represent ±standard deviation (±SD). Unpaired Student’s t-test was used to compare data from two groups. Pearson correlation coefficients were calculated using the Coloc2 plugin in ImageJ. Plots were made in Matlab or Prism.

## Acknowledgments

We thank Wenjing Sun for sharing nucleosome arrays, Dr. Linyu Zuo from Dr. Zhi Qi’s lab at Peking University for sharing original luciferase assay plasmids, the Tsinghua Nikon imaging center for technical assistance, other members in the Li Lab for discussion, and the support of Tsinghua Xuetang Life Science Program. This work was supported by grants from the National Key R&D Program (2019YFA0508403 to P.L.) and the Natural Science Foundation of China (32150023, 32125010 to P.L.).

## Author contributions

G.L. and P.L. conceived the project. G.L. performed all experiments and data analyses. P.L. supervised the experiments; G.L. and P.L. wrote the paper.

## Data and materials availability

All data are available in the main text or the supplementary materials. Expression plasmids and cell lines are available upon request.

