## Supplementary Information for "PHF1 compartmentalizes PRC2 at target loci via phase separation"

Genzhe Lu[1, 2, 3, 4, #] and Pilong Li[1, 2, #]

1 Beijing Frontier Research Center for Biological Structure, School of Life Sciences, Tsinghua University, Beijing 100084, China

2 Tsinghua-Peking Joint Center for Life Sciences, Beijing 100084, China

3 Tsinghua Xuetang Life Science Program, Tsinghua University, Beijing 100084, China

4 Present Address: The Rockefeller University, New York, NY 10065, USA

Supplementary information

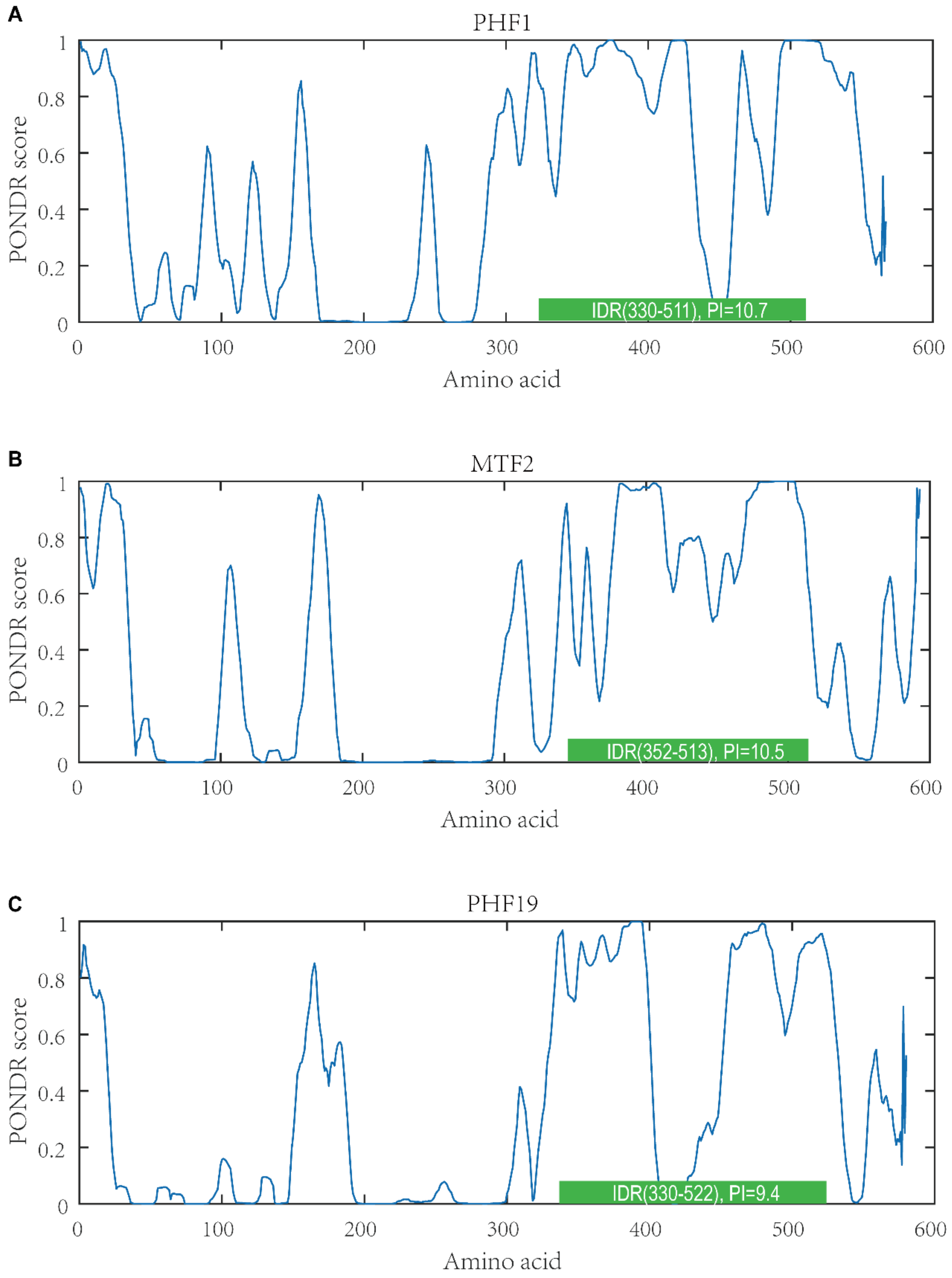


**Figure S1. IDR predictions of PCL proteins.** Plots of PONDR scores with respect to amino acid numbers of PHF1 (**A**), MTF2 (**B**), and PHF19 (**C**). IDR regions and isoelectric points of IDRs are indicated in green boxes.

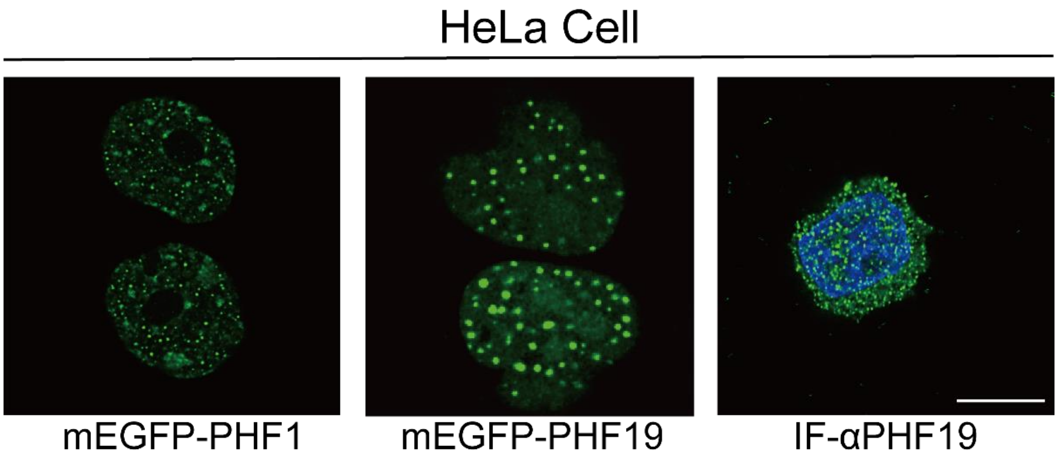


**Figure S2.** Confocal images of mEGFP-PHF1 and mEGFP-PHF19 in living cells, and immunofluorescence staining of PHF19 (IF-αPHF19) in HeLa cells. Nuclei were stained by DAPI. Scale bar, 10 μm. Related to Figure 1.

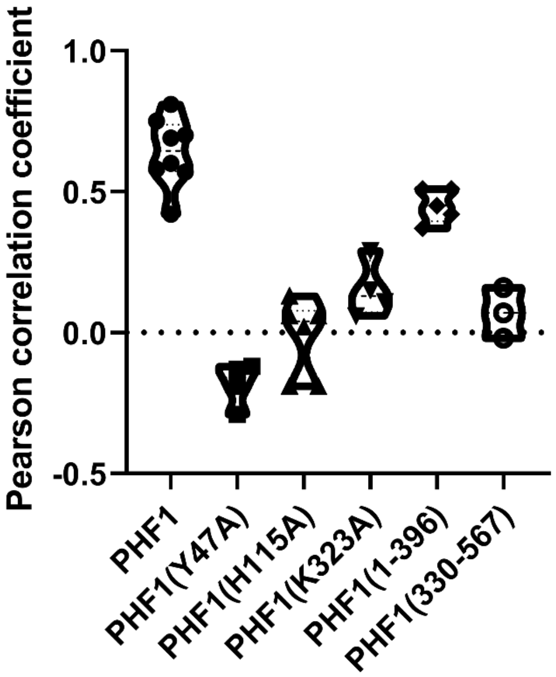


**Figure S3. Pearson correlation coefficients for colocalization of PHF1 variants with H3K27me3, related to Figure 2.**

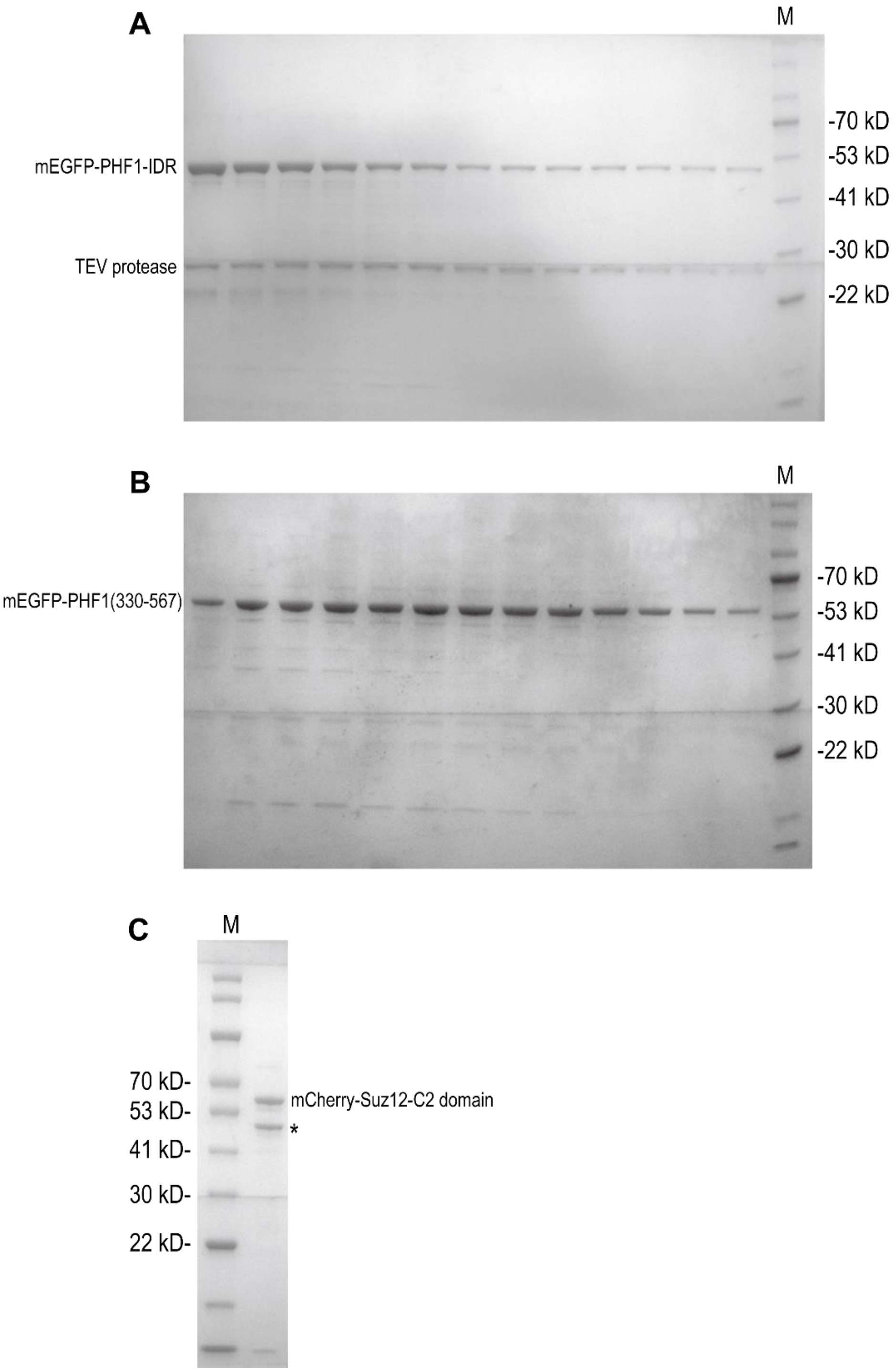


**Figure S4. SDS-PAGE gels of purified proteins. A** Gel filtration peak fractions of mEGFP-PHF1-IDR. **B** Gel filtration peak fractions of mEGFP-PHF1(330-567). **C** mCherry-Suz12-C2 domain. * denotes a degradation band.

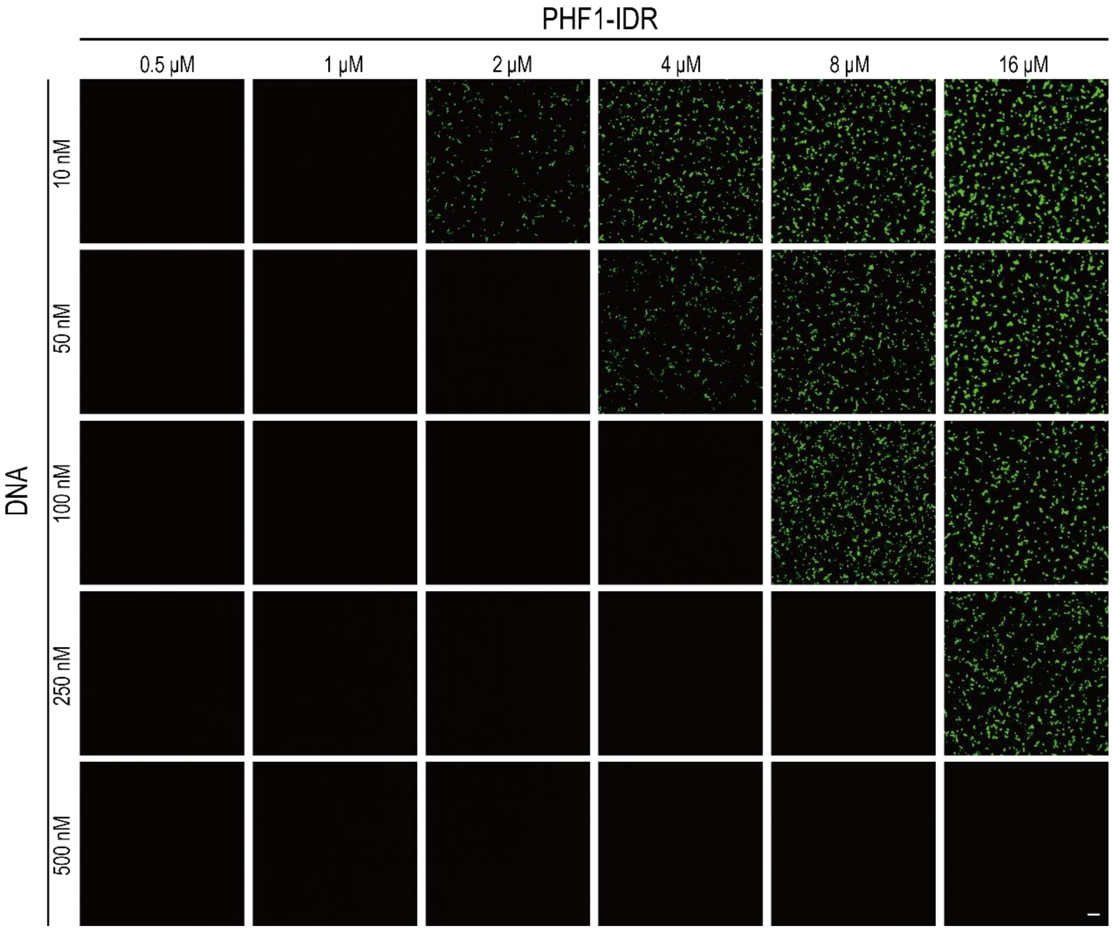


**Figure S5. Complete phase diagram of PHF1-IDR with 72 bp DNA, related to Figure 3.** Scale bar, 10 μm

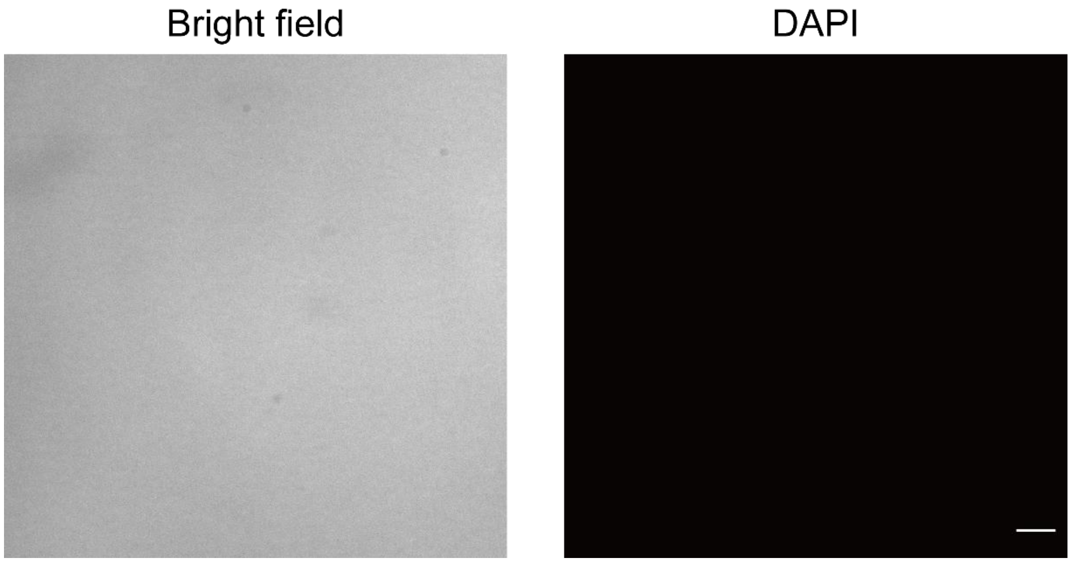


**Figure S6.185 nM 12X H3K27me3 nucleosome arrays do not phase separate at 100 mM NaCl, related to Figure 3F.**

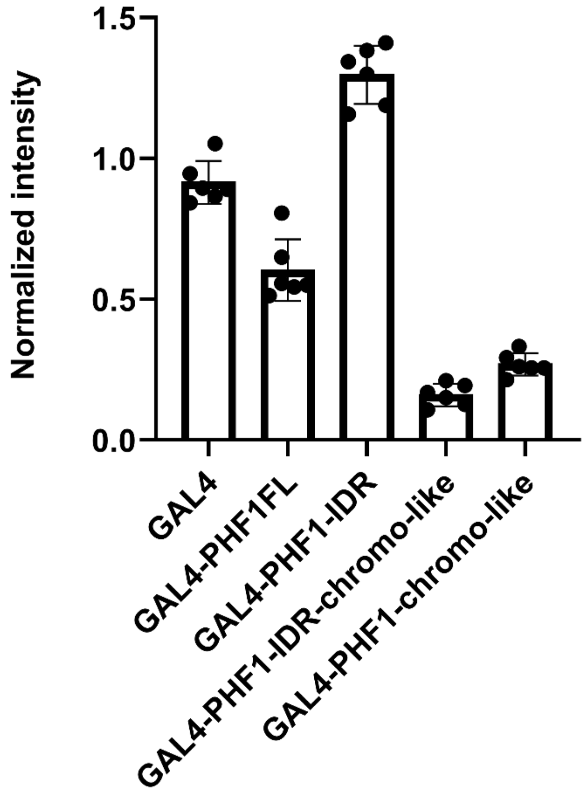


**Figure S7. Complete luciferase reporter assay results (n=6), related to Figure 4.** Data for GAL4-PHF1-IDR-chromo-like and GAL4-PHF1-chromo-like are the same as Figure 4B and are shown here for better comparison.
